# Sexual maturation in Atlantic salmon male parr may be triggered both in early spring and late summer under standard farming conditions

**DOI:** 10.1101/2021.03.29.437499

**Authors:** Elia Ciani, Kristine von Krogh, Rasoul Nourizadeh-Lillabadi, Ian Mayer, Romain Fontaine, Finn-Arne Weltzien

## Abstract

Male Atlantic salmon (*Salmo salar*) display different sexual strategies, maturing either as parr during the freshwater phase (as sneaky spawners), or as post smolts following one or several years at sea. First sexual maturation (puberty) occurs at different times depending on environmental and genetic factors. To improve our knowledge on the timing (age and season) of first sexual maturation in Atlantic salmon male parr, we investigated pubertal activation in second generation farmed salmon from the Norwegian river Figgjo, reared under natural conditions of photoperiod and water temperature. Histological analysis, in combination with morphometric measurements, plasma androgen levels and pituitary gonadotropin gene expression analysis revealed that, as previously reported, some male parr initiated early sexual maturation in spring at one year of age. Interestingly, some male parr were observed to initiate sexual maturation already in autumn, six months after hatching (under-yearlings), much earlier than reported in previous studies. One-year old maturing males showed a low induction in gonadotropin levels, while under-yearling maturing males displayed a significant increase in *fshb* transcripts as compared to immature fish. Plasma testosterone, detectable also in immature males, increased constantly during testes development, while 11-ketotestosterone, undetectable in immature and early maturing males, increased during more advanced stages of maturation. A mild feminization of the testes (ovotestes) was detected in a subset of samples. This study brings new knowledge on the little investigated field of sexually maturing under-yearlings in Atlantic salmon. This is also the first study comparing the physiology of under-yearling vs one-year old maturing male parr, thus bringing new insights to the remarkable plasticity of Atlantic salmon puberty.

## 1 Introduction

Puberty is a process of physical and physiological changes through which an individual reaches sexual maturation for the first time in life. In male teleosts, as in other male vertebrates, puberty is characterized by the onset of spermatogenesis (Schulz and Miura, 2002). Spermatogenesis is regulated by the brain-pituitary-gonadal (BPG) axis, with the pituitary producing the gonadotropins, follicle-stimulating hormone (Fsh) and luteinizing hormone (Lh), which are key hormones involved in the control of gonadal maturation (Levavi-Sivan et al., 2010).

Fsh and Lh are heterodimeric glycoproteins consisting of two non-covalently linked subunits, a common α-(Gpα), and a hormone-specific β-subunit (Lhβ or Fshβ) that confers the biological activity (Pierce and Parsons, 1981; Swanson et al., 2003). Unlike mammals, where both gonadotropins are produced from the same pituitary cell, Fsh and Lh in teleosts are mostly produced from two different cell types, located in the *proximal pars distalis* (PPD) of the pituitary gland (Fontaine et al., 2020a; Levavi-Sivan et al., 2010; Weltzien et al., 2004).

The physiological role of gonadotropins has been extensively studied in salmonids (Gomez et al., 1999; Maugars and Schmitz, 2008, 2006; Prat et al., 1996; Swanson et al., 1989). In males, Fsh and Lh induce the production of the androgens 11-ketotestosterone (11-KT) and testosterone (T) with similar efficacy during early stages of maturation (Planas et al., 1993; Planas and Swanson, 1995). 11-KT, which is the main androgen in salmonids (Borg, 1994; Rege et al., 2019), plays a major role in all stages of teleost spermatogenesis, being involved in the induction of spermatogonial proliferation, meiotic division and spermiogenesis (Miura et al., 1991). T is involved in several feedback mechanisms to the hypothalamus (Amano et al., 1994; Fontaine et al., 2020b; Goos et al., 1986) and pituitary (Fontaine et al., 2020b; Montero et al., 1995; Xiong et al., 1993), and can act through both the androgen receptor (Ar) and the estrogen receptors (Ers). In addition, Lh is a potent stimulator of the maturation-inducing steroid 17α,20β-dihydroxy-4-pregnen-3-one (DHP) during the last stages of maturation in both sexes (Planas and Swanson, 1995).

In salmonids, plasma Fsh levels increase already during the onset of spermatogenesis, earlier than Lh plasma levels, which remain undetectable or very low during the initial stages of maturation and rises only during spermiation (Swanson et al., 2003). It has been suggested that Fsh, due to its presence in the plasma of immature fish and its capacity to stimulate both steroidogenesis and spermatogonial proliferation (Loir, 1999), plays a major role during the early stages of gonadal development in salmonids, while Lh is more involved in the later stages of maturation (Levavi-Sivan et al., 2010; Schulz et al., 2010; Yaron et al., 2003).

The anadromous Atlantic salmon (*Salmo salar*) shows remarkable plasticity concerning male sexual maturation. Males may mature either during the freshwater phase (parr), from one to four years of age, as small “sneaky spawners/precocious males” (Aas et al., 2011), or in the saltwater phase following smoltification at a larger size as one-(grilse) or multi-sea-winter individuals (Garcia De Leaniz et al., 2007; Hutchings and Jones, 1998; Taylor, 1991). There has also been occurrences of male parr maturation earlier than one year of age (under-yearling or 0+) both in wild (Bagliniere and Maisse, 1985) and farmed populations reared under artificial light regimes (Nordgarden et al., 2007). However, little is known about the physiology of under-yearling maturing males regarding gonadotropin expression, testes development or morphometric profiles. Here, we recorded the timing of seasonal sexual maturation in farmed Atlantic salmon male parr reared in south-west Norway under natural conditions of photoperiod and water temperature. The aim was to assess the specific time and age at which sexual maturation is initiated and to compare the physiology of fish maturing as under-yearling or as one-year old individuals.

## 2 Material and Methods

### 2.1 Animals and rearing conditions

One-year old Atlantic salmon male parr were sampled in spring/summer 2017 (batch 1), in addition to a batch of under-yearling (four months old) parr in summer/autumn 2017 (batch 2). The broodstock was a farmed first generation from wild caught fish in the river Figgjo (south-west Norway, 58°47’ N 5°47’ E). The fish were reared at the Norwegian Institute for Nature Research (NINA) salmon research station at Ims, Norway (58°54’N, 5°57’E), hatching on February 22^nd^, 2016 (batch 1), and March 14^th^, 2017 (batch 2). After first feeding, fish were reared in outdoor tanks (volume 7.8 m^3^), under natural conditions regarding photoperiod and temperature (Supplementary figures S1, S2). All experiments were performed in accordance with guidelines of the Animal Welfare Committee of the Norwegian university of life sciences (NMBU) and of the European Union regulation concerning the protection of experimental animals (Directive 2010/63/EU). Appropriate measures were taken during sampling to minimize pain and discomfort (FOTS application ID12523).

### 2.2 Sampling

Fish were anesthetized with MS222 (80 mg/l; Pharmaq, Overhalla, Norway) and euthanized via quick decapitation. Body weight, gonad weight and fork length were measured for all fish. Gonadosomatic index (GSI = gonad weight / body weight x 100) and condition factor (K = 100 body weight / fork lenght^3^) were calculated from the morphometric measurements. As testes size increases at puberty, the gonad somatic index was used to initially divide males as either immature or maturing before additional analysis. The fish were divided into two groups based on GSI and morphology: one group with immature thread-like testes (GSI < 0.05), and a second group showing the first signs of testicular development (GSI > 0.05). Without a universal criterion within the literature for a threshold GSI value to classify males as immature (Duston and Saunders, 1999; Endal et al., 2000; Kadri et al., 1997), we initially classified males with GSI ≤ 0.05 as immature while those with GSI > 0.05 as maturing. This threshold is also supported by our histology results as shown below. Fish were sampled until a minimum of six biological replicates per group were obtained (Table 1, 2). One year old parr (14 months old at first sampling) were sampled from April to July 2017 (25^th^ April; 8^th^ and 23^rd^ May; 7^th^ June; 4^th^ July) while under-yearling parr (4 months old at first sampling) were sampled from July to October 2017 (4^th^ July; 22^nd^ August; 27^th^ September; 23^rd^ October).

**Table 1.** Summary of morphometric measurement, steroid plasma levels and relative mRNA abundance in one-year old male parr measured in the present study. mRNA levels are represented as fold induction to the lowest point. (K) condition factor (m) maturing (i) immature (SD) standard deviation (ND) not detected (NA) not available.

|  |  |  | 25 Apr | 8 May | 23 May | 7 Jun | 4 Jul |
| --- | --- | --- | --- | --- | --- | --- | --- |
| Fork length (cm) | m | mean | 10.63 | 10.01 | 11.57 | 12.72 | 13.55 |
|  |  | SEM | 0.23 | 0.35 | 0.40 | 0.35 | 0.33 |
|  |  | replicates | 7 | 7 | 7 | 12 | 6 |
|  | i | mean | 9.23 | 9.07 | 9.67 | 10.70 | NA |
|  |  | SEM | 0.37 | 0.27 | 0.31 | - |  |
|  |  | replicates | 7 | 6 | 6 | 1 | 0 |
| Body weight (cm) | m | mean | 12.62 | 11.46 | 18.67 | 26.10 | 31.98 |
|  |  | SEM | 1.33 | 1.13 | 2.34 | 2.62 | 2.42 |
|  |  | replicates | 7 | 8 | 7 | 12 | 6 |
|  | i | mean | 7.82 | 7.37 | 9.10 | 10.90 | NA |
|  |  | SEM | 0.89 | 0.58 | 1.21 | - |  |
|  |  | replicates | 7 | 6 | 6 | 1 | 0 |
| GSI | m | mean | 0.236 | 0.392 | 0.375 | 0.401 | 0.852 |
|  |  | SEM | 0.064 | 0.142 | 0.099 | 0.048 | 0.132 |
|  |  | replicates | 7 | 8 | 7 | 12 | 6 |
|  | i | mean | 0.042 | 0.040 | 0.021 | 0.037 | NA |
|  |  | SEM | 0.004 | 0.007 | 0.005 | - |  |
|  |  | replicates | 7 | 6 | 6 | 1 | 0 |
| K | m | mean | 1.03 | 1.06 | 1.15 | 1.21 | 1.27 |
|  |  | SEM | 0.05 | 0.03 | 0.04 | 0.04 | 0.02 |
|  |  | replicates | 7 | 7 | 7 | 12 | 6 |
|  | i | mean | 0.98 | 0.98 | 0.97 | 0.89 | NA |
|  |  | SEM | 0.05 | 0.03 | 0.05 | - |  |
|  |  | replicates | 7 | 6 | 6 | 1 | 0 |
| 11KT (ng/μl) | m | mean | 0.09 | 0.13 | 0.23 | 0.46 | 2.07 |
|  |  | SEM | 0.09 | 0.08 | 0.14 | 0.11 | 0.57 |
|  |  | replicates | 7 | 7 | 7 | 12 | 6 |
|  | i | mean | ND | ND | ND | ND | NA |
|  |  | SEM |  |  |  |  |  |
|  |  | replicates | 7 | 5 | 6 | 1 | 0 |
| T (ng/μl) | m | mean | 0.74 | 0.96 | 1.11 | 1.27 | 1.72 |
|  |  | SEM | 0.07 | 0.10 | 0.09 | 0.14 | 0.16 |
|  |  | replicates | 6 | 7 | 7 | 12 | 6 |
|  | i | mean | 0.35 | 0.22 | 0.34 | NA | NA |
|  |  | SEM | 0.15 | 0.13 | 0.10 |  |  |
|  |  | replicates | 6 | 5 | 4 | 0 | 0 |

**Table 2.** Summary of morphometric measurement, steroid plasma levels and relative mRNA abundance in under-yearling male parr measured in the present study. mRNA levels are represented as fold induction to the lowest point. (K) condition factor (m) maturing (i) immature (SD) standard deviation (ND) not detected (NA) not available.

|  |  |  | 4-Jul | 22-Aug | 27-Sep | 23-Oct |
| --- | --- | --- | --- | --- | --- | --- |
| Fork length cm | m | mean |  |  | 9,48 | 10,00 |
|  |  | SEM |  |  | 0,33 | 0,31 |
|  |  | replicates |  |  | 5 | 9 |
|  | i | mean | 6,00 | 9,50 | 11,32 | 9,91 |
|  |  | SEM | 0,18 | 0,25 | 0,32 | 0,60 |
|  |  | replicates | 5 | 6 | 9 | 8 |
| Body weight cm | m | mean |  |  | 8,92 | 10,66 |
|  |  | SEM |  |  | 0,87 | 1,10 |
|  |  | replicates |  |  | 5 | 9 |
|  | i | mean | 2,34 | 9,70 | 16,19 | 12,00 |
|  |  | SEM | 0,21 | 0,93 | 1,34 | 0,95 |
|  |  | replicates | 5 | 6 | 9 | 8 |
| GSI | m | mean |  |  | 0,168 | 1,710 |
|  |  | SEM |  |  | 0,059 | 0,945 |
|  |  | replicates |  |  | 5 | 9 |
|  | i | mean | 0,010 | 0,026 | 0,034 | 0,034 |
|  |  | SEM | 0,003 | 0,002 | 0,003 | 0,005 |
|  |  | replicates | 5 | 6 | 9 | 8 |
| K | m | mean |  |  | 1,03 | 1,04 |
|  |  | SEM |  |  | 0,02 | 0,02 |
|  |  | replicates |  |  | 5 | 9 |
|  | i | mean | 1,07 | 1,11 | 1,09 | 1,50 |
|  |  | SEM | 0,05 | 0,02 | 0,02 | 0,41 |
|  |  | replicates | 5 | 6 | 8 | 8 |

### 2.3 Testis histology

Testes from all sampled fish were dissected and fixated in 4 % glutaraldehyde (VWR, Radnor, USA) overnight at 4 °C. The samples where then stored in 70 % ethanol (EtOH) at 4 °C until further processing. Histological analyses were performed as follows: tissues were dehydrated with EtOH washes at increasing concentrations (up to 100 % EtOH), each lasting 30 min. The last step was repeated three times. Tissues were then kept at room temperature overnight slowly shaking in preparation solution (100 ml Technovit 7100 added 1 g of Hardener I (Heraeus Kulzer, Hanau, Germany)). Afterwards, tissues were embedded in cold Histoform S (Heraeus Kulzer) added approx. one mL preparation solution with 50 μL Hardener II (Heraeus Kulzer) and incubated at 37 °C. Finally, samples were mounted on Histoblocs using Technovit 3040 (both from Heraeus Kulzer).

Sagittal (for small and medium testes) and transverse (for large testes) sections, 3 μm thick, were prepared using a Leica RM2245 microtome (Leica Biosystems, Wetzlar, Germany). Sections were separated by at least 30 μm and collected from the periphery until the middle of the tissue. Dried sections were stained with Toluidine Blue O (Sigma-Aldrich) and mounted with Coverquick 4000 (VWR International, Radnor, PA, USA) before histological analysis. The maturational stages of the testes were determined by the most advanced germ cell present in the tissue (Table 4). Germ cell stages were defined according to the description by Melo et al. (2014). Histological analyses were performed in all maturing and two randomly selected immature fish per time point. At least five sections per testes were analysed.

**Table 4.** Stages of testes development used for histological analysis. Each stage is defined by the most developed germ cell cyst present in the testis tissue. SPA, spermatogonia A, either undifferentiated or differentiated; SPB, spermatogonia B; SC, spermatocytes; ST, spermatids; SZ, spermatozoa.

| Stages | Observed cysts |
| --- | --- |
| I | SPA |
| II | SPA+SPB |
| III | SPA+SPB+SC |
| IV | SPA+SPB+SC+ST |
| V | SPA+SPB+SC+ST+SZ<br>(With some tubules still immature, or early maturing) |
| VI | SZ is dominating. Large lumen in tubules |

### 2.4 Gene expression analysis

Individual pituitaries from at least six maturing and six immature fish per time point were collected and stored in 300 μl TRIzol reagent (Invitrogen, Waltham, USA) overnight at 4 °C, then at −20 °C. Total RNA was isolated using TRIzol reagent according to the manufacturer’s instructions. To avoid any genomic contamination, extracted RNA was treated with 2 U DNase (TURBO DNA-free kit, Ambion, Foster City, USA). The concentration of total RNA was measured via Qubit Fluorometer (Invitrogen) using Qubit RNA BR Assay Kit (Invitrogen). The 260/280 ratio for all samples was between 1.8 and 2. Quality of RNA samples was measured using Bioanalyzer 2100 (Agilent, Santa Clara, USA). All samples showed RNA Integrity Number (RIN) above 8. SuperScript III reverse transcriptase (Invitrogen) and 5 μM random hexamer primers (Invitrogen) were used, following producer’s instructions, to reverse transcribe 170 ng total RNA from individual pituitaries.

Primer-Blast from NCBI website (Ye et al., 2012) was used to design specific qPCR primer sets for the genes of interest (Table 3). Primers were tested for primer dimers and hairpin potential with Vector NTI Express software (Lu and Moriyama, 2004). Gene expression of the target genes *fshb* and *lhb* was quantified by qPCR using the Light Cycler 96 (Roche, Basel, Switzerland) thermocycler and SYBR Green I master (Roche) kit as shown in Weltzien et al. (2005). In brief, each individual sample was run in duplicate using 3 μL of the same cDNA dilution 1/10. A negative control and an inter-plate calibrator were present in triplicate for each qPCR plate. Real-time conditions were 10 min incubation at 95 °C followed by 40 cycles at 95 °C for 10 s, 60 °C for 10 s, and 72 °C for 8 s. The specificity of the amplified product was verified by running a melting curve analysis and by sequencing of the PCR product. Standard curves were run in triplicates using dilution series obtained from RT products from pooled total RNA of pituitaries. The relative abundance of transcript was determined using GenEx software (Mangalam et al., 2001) through the algorithms outlined by Vandesompele et al. (2002). The reference genes used for data normalization were *rna18s* and *ef1a*. Stability of the reference genes was tested using the online tool RefFinder (Kim et al., 2010).

**Table 3.**
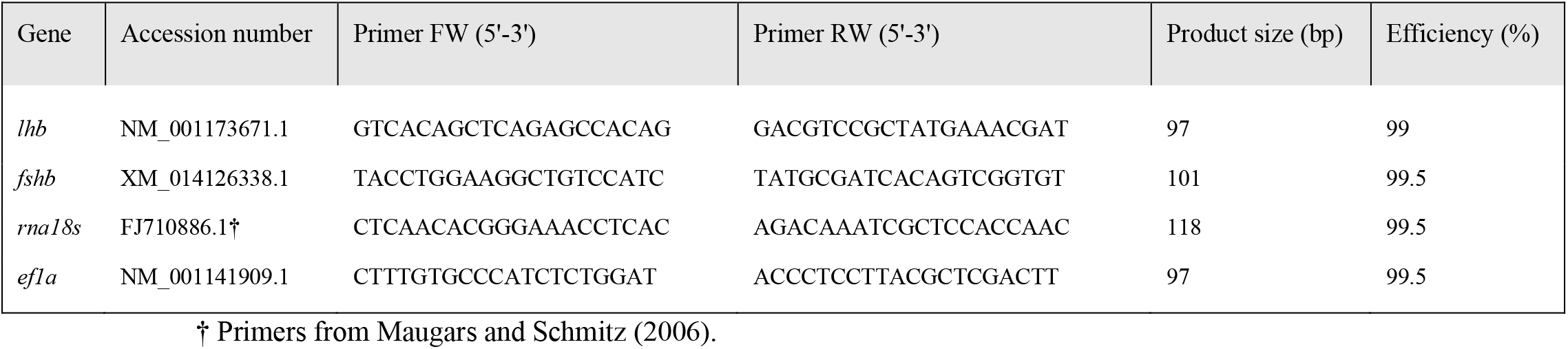
Primers used in the present study.

### 2.5 Radioimmunoassay (RIA)

Blood from all sampled fish was collected from the caudal vein with heparinised (Sigma-Aldrich) syringes. Plasma was then isolated by centrifugation and stored at −80 °C. Levels of T and 11-KT were measured by specific radioimmunoassay (RIA) as previously described by Mayer et al. (1990). Standard curves were prepared mixing steroid solutions containing decreasing concentrations of the respective, non-radioactive steroid in pH 7.0 RIA buffer (NaH_2_PO_4_ 3.87 g, Na_2_HPO_4_ 10.67 g, Na-Azid 0.05 g, NaCl 9 g, gelatine 1 g, ddH_2_O 1 L). The inter-assay variations for T and 11-KT were 13.5 and 14 %, while the intra-assay variations were 5.6 and 4.8 %, respectively. The analysis of 11-KT plasma levels were prioritised when the amount of plasma isolated was not sufficient for both analysis. Due to the small size of the fish and the insufficient amount of plasma available, RIA was not performed on underyearling fish (Table 1, 2)

### 2.6 Statistics

All statistical analyses were performed with JMP pro V14.1 software (SAS Institute Inc., Cary, NC, USA). Shapiro-Wilk W test was used for testing normality and Quantile Range method (Q=3; Tail 0.1) was used to identify and exclude outliers. When needed, data were log transformed to meet test criteria. Variations in morphometric data and gene expression during sexual maturation were analysed for statistical significance by two-way ANOVA, followed by Tukey’s HSD test. Variations in gene expression relative to testes developmental stage were assessed using one-way ANOVA, followed by Tukey’s HSD test. In this study, p values <0.05 were considered statistically significant.

## 3 Results

All results of the present study are summarised in Table 1 and Table 2. All values are reported in the following chapter as mean ± SEM.

### 3.1 Testes histology

Examples of the testes developmental stages identified in the present study (I – VI), are presented in Fig. 1 (A-F). All fish classified as immature according to GSI ≤ 0.05, had stage I testis, indicating that active spermatogenesis had not started, and only spermatogonia type A cells were present in the tissue. In contrast, in all males with GSI > 0.05, testis development had progressed into stage II to IV, indicating that active spermatogenesis had started, with spermatogonia type B cells, spermatocytes and spermatids found in the testicular lobules. This classification, based on GSI, was used to categorize between non- and maturing fish.

**Figure 1.**
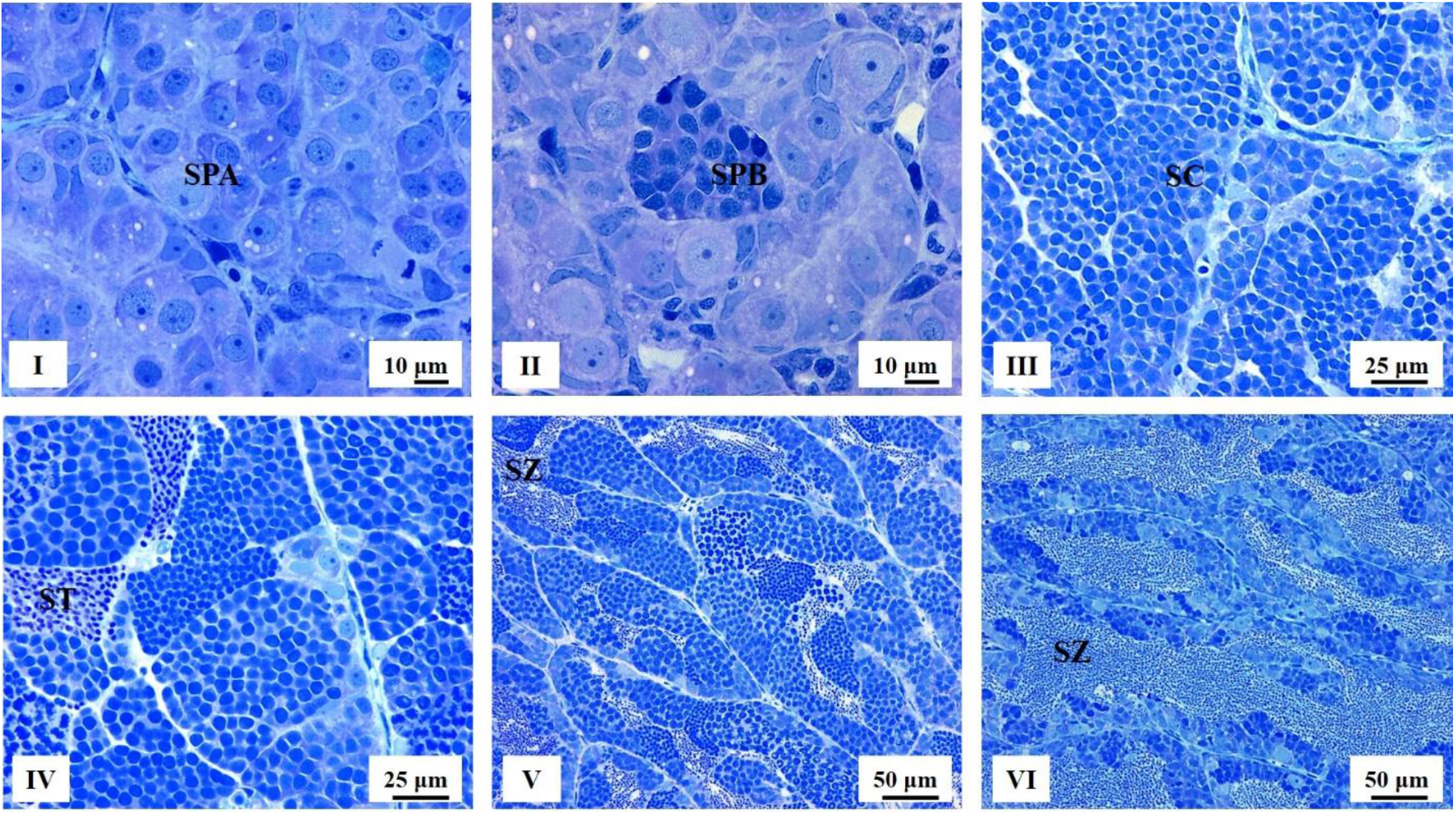
Testis developmental stages in Atlantic salmon parr defined according to the most advanced cyst present: Stage **I**, spermatogonia A (SPA) predominate, either undifferentiated or differentiated; Stage **II**, occurrence of spermatogonia B (SPB); Stage **III**, spermatocytes (SC) present; Stage **IV**, spermatids (ST) present; Stage **V**, first signs of spermatozoa (SZ), with some tubules still immature, or early maturing. Stage **VI**, spermatozoa dominate in all tubules. Large lumen in tubules. Sections (3 μm) were prepared in plastic resin and stained with Toluidine Blue O.

In one-year old maturing fish, the testes development had reached stage II already by the first sampling on April 25^th^ (Fig 2A). On May 8^th^, stage II and III testes were predominantly detected, although a few individual had developed into stage IV, V and VI. The occurrence of the most advanced stages (IV to VI) increased during the season, and on July 4^th^, only stage V and VI were detected in the examined tissue (Fig 2A). Of note, histologically, a stage IV testis with residual spermatozoa is not distinguishable from a stage V. Similarly, as stage V and VI include spermatozoa, we did not distinguish residual from new spermatozoa. Interestingly, the presence of residual spermatozoa (Fig 3A) was observed in 6 out of 27 (22.2%) testes at stages I, II and III, lacking one or more intermediate germ cell stages suggesting that complete maturation has already performed by those fish previously. Thus in the rest of the study, we have investigated both one-year old and under-yearling fish.

**Figure 2.**
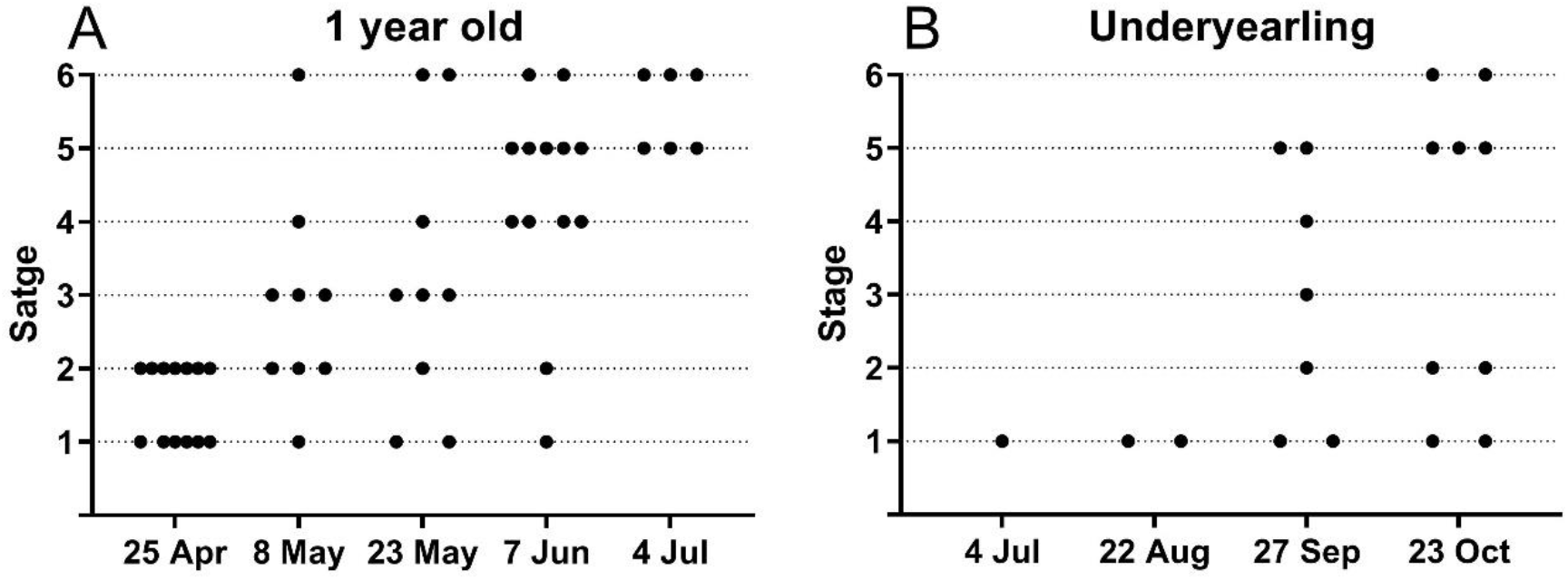
Advancement in testis developmental stages in **A)** one-year old and **B)** under-yearling male parr in spring/autumn. Each dot represents the testis developmental stage assigned according to the most developed cyst detected. Testes of all maturing and a few randomly selected non-maturing males were analysed at each date.

**Figure 3.**
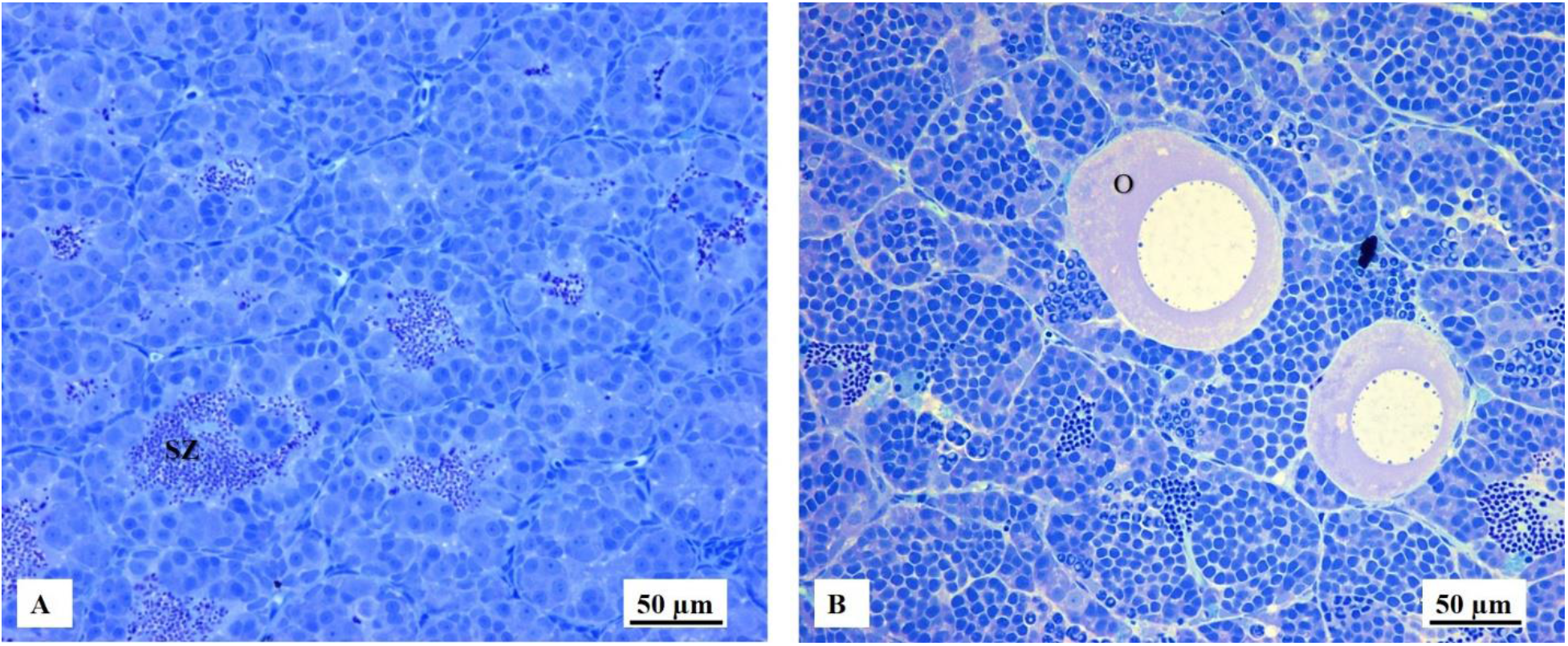
Sections of testes showing **A)** residual spermatozoa (SZ) and **B)** early previtellogenic oocytes (O). Sections (3 μm) were prepared in plastic resin and stained with Toluidine Blue O.

In under-yearling males, the first signs of testis maturation beyond proliferating SPA were detected on September 27^th^, with testes stages spanning from II to V. By October 23^rd^, all examined samples had reached full maturation (stage V and VI) (Fig 2B).

There were some peculiarities detected in the histological analysis. First, the presence of previtellogenic oocytes within the testis tissue (Fig 3B) was observed in 6 out of 69 (8.7%) analysed testes, and this occurred in both under-yearlings and one-year old fish. It is possible that the absolute number of fish with testis-ova may be higher as we did not analyse all the tissue from each testis, and furthermore, that oogonia and peri-nucleolar oocytes can be difficult to distinguish from undifferentiated spermatogonia Type A by histology alone. Second, in several testes from both one-year old and under-yearling maturing males, an incomplete tissue development with part of the testes fully developed and others being completely immature, was detected (Supplementary figure S3).

### 3.2 Biometry

One-year old maturing fish (GSI > 0.5) were already identified from the first sampling on April 25^th^ (Fig 4A). Maturing and immature fish display no significant differences in fork length (Fig 4B) and body weight (Fig 4C) except on May 23^rd^, where mature fish have increased fork length and body weight. No differences between groups at any time point are detected for condition factor (Fig 4D). Due to the lower number of parr available in the batch and the high percentage of maturing fish within the parr population, only one immature individual is present on June 7^th^. Data from this fish are shown in figure 1 for graphical purposes only but are not included in the statistical analysis. All fish sampled on July 4^th^ were classified as maturing.

**Figure 4.**
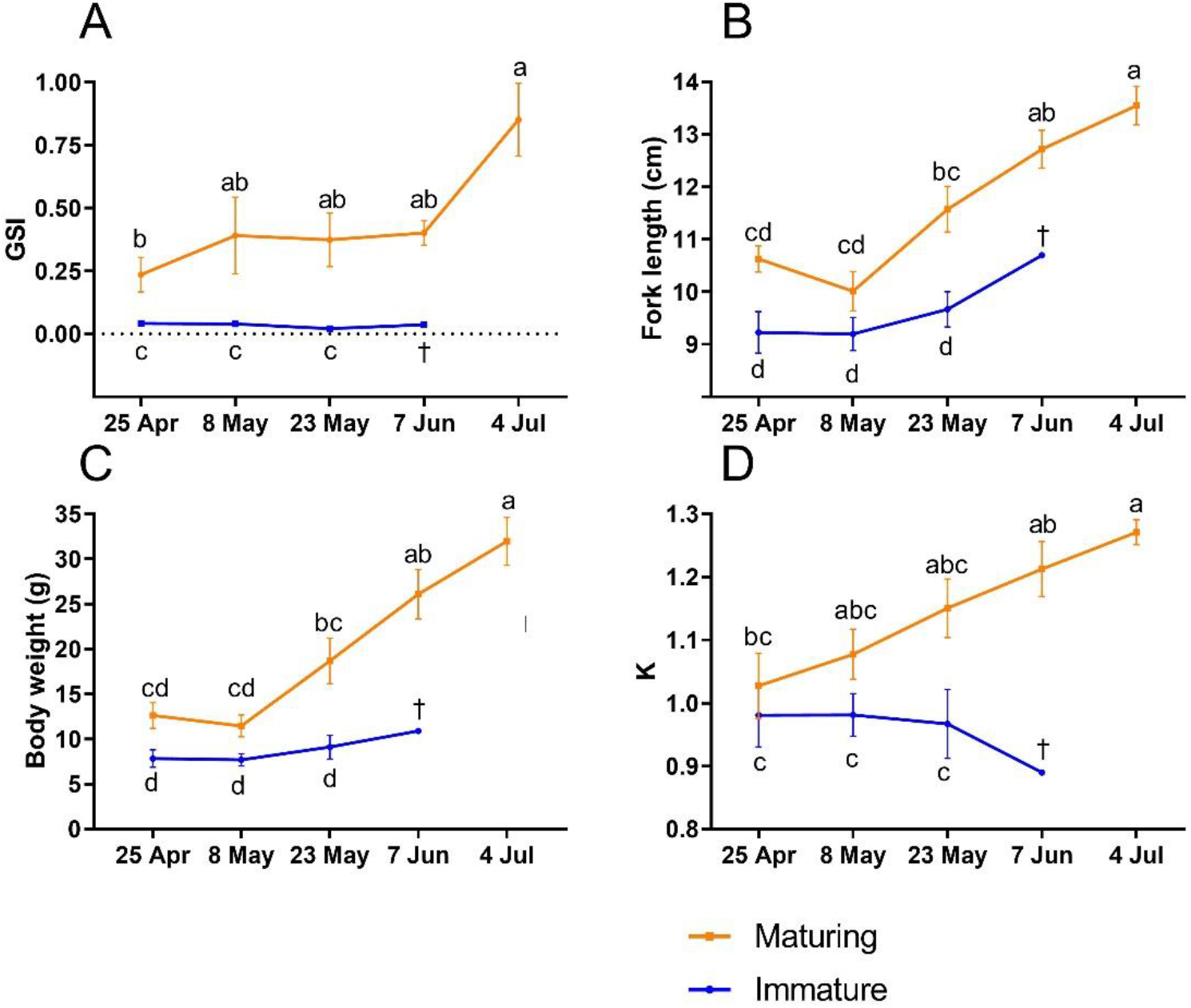
Morphometric measurements of one-year old male parr during spring/summer. **A)** gonadosomatic index (GSI = gonad weight/body weight*100); **B)** Fork length (L); **C)** body weight (W); **D)** condition factor (K=100 W/L^3^). Data are presented as mean ± SEM. Distinct letters denote statistically significant differences among groups (p<0.05), analysed via two-way ANOVA followed by Tukey multiple comparison test. † Only one immature fish was available on June 7^th^. It is presented for graphical purposes only and is not included in the statistical analysis. The number of biological replicates per data point is listed in table 1.

Under-yearling maturing fish (GSI > 0.5) were detectable from September 27^th^, six months after hatching (Fig 5A). No statistically significant differences were measured between groups in fork length at any time point (Fig 5B), while immature fish displayed greater body weight compared to maturing fish on September 27^th^ (Fig 5C). No differences in condition factor were detected between groups (Fig 5 D).

**Figure 5.**
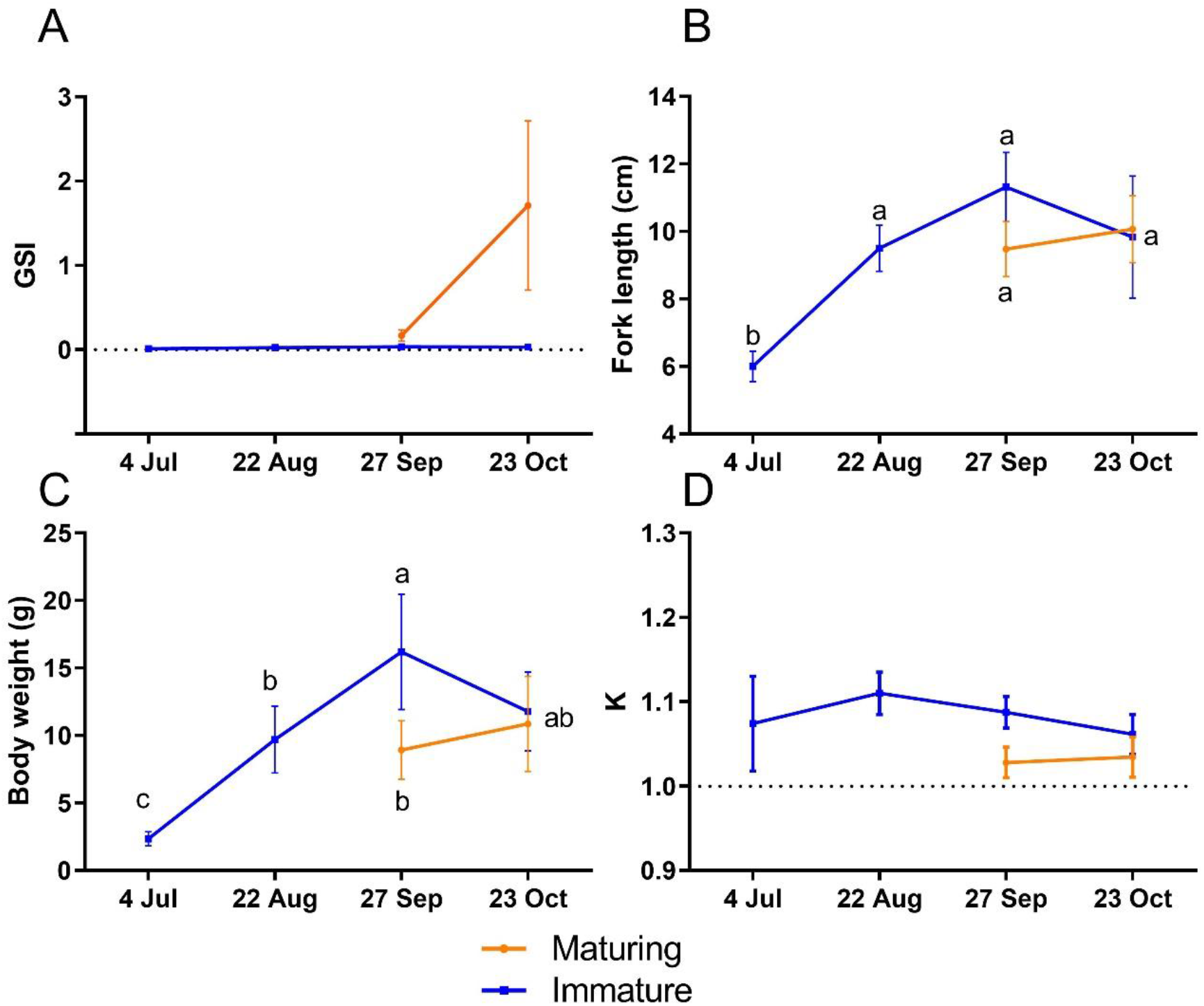
Morphometric measurements of under-yearling male parr during summer/autumn. **A)** Gonadosomatic index (GSI= gonad weight/body weight*100); **B)** Fork length (L); **C)** Body weight (W); **D)** Condition factor (K=100 W/L^3^); Data are shown as mean ± SEM. Distinct letters denote statistically significant differences among groups (p<0.05), analysed via two-way ANOVA followed by Tukey multiple comparison test. The number of biological replicates per point is listed in table 2.

### 3.3 Seasonal gonadotropin expression

In one year old maturing males pituitary *fshb* mRNA levels remained stable over time (from April 25^th^ to July 4^th^) while they decreased in immature fish from April 25^th^ to May 23^rd^. On May 23^rd^, maturing fish displayed higher *fshb* transcripts than immature fish. (Fig 6A). Pituitary *lhb* mRNA transcripts decreased over time in both maturing and immature fish from April to May/June. Significantly higher *lhb* mRNA levels are detected in maturing fish as compared to in immature fish only on May 23^rd^ (Fig 6C).

**Figure 6.**
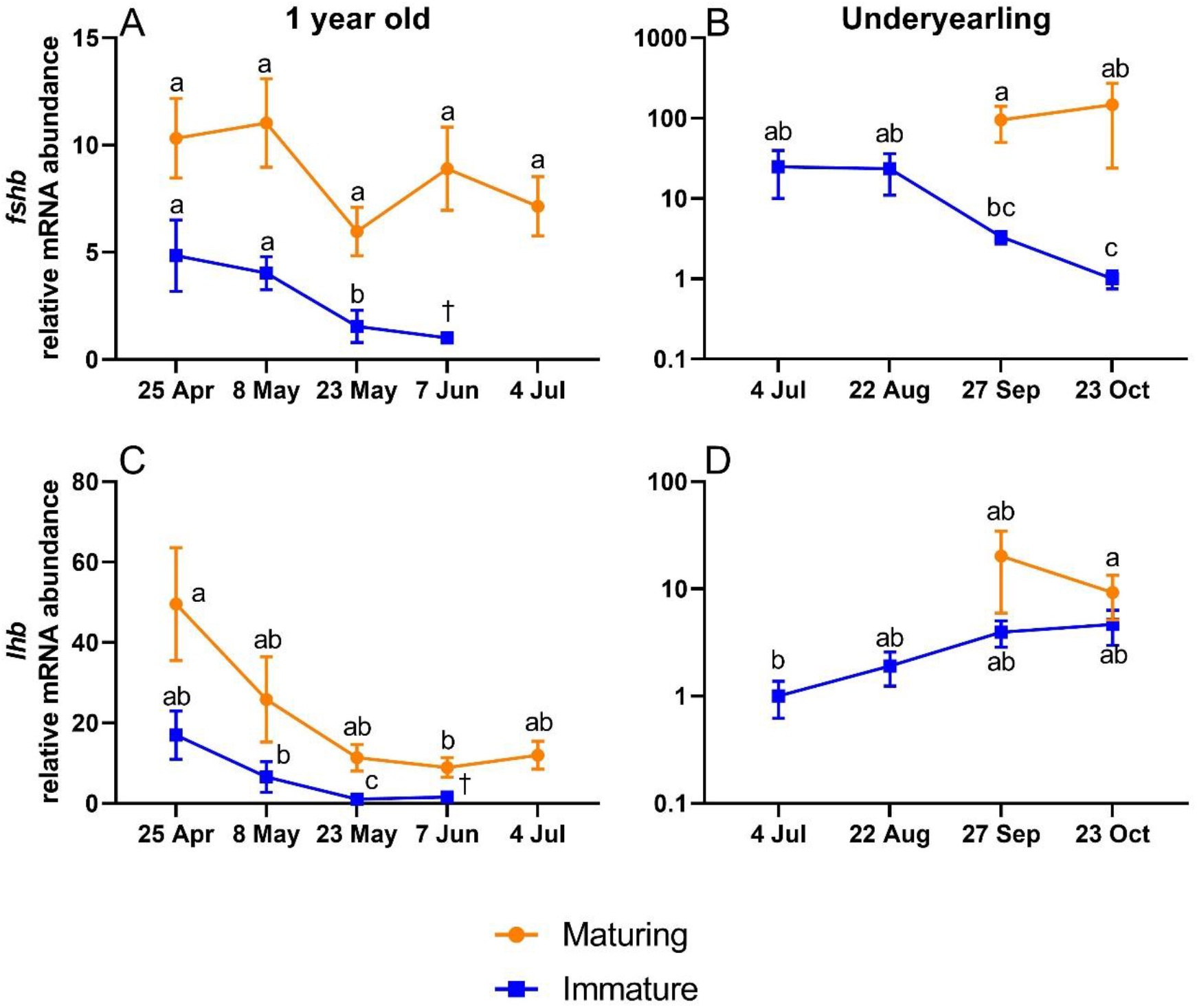
Relative abundance of: **A-B)** *fshb* and **C-D)** *lhb* mRNA in pituitaries from **A-C)** one-year old and **B-D)** under-yearling male Atlantic salmon parr. Data are graphically presented as fold change of the lowest expressing point (set as value 1) (mean ± SEM; n=6). Distinct letters denote statistically significant differences among groups (p<0.05), analysed via two-way ANOVA followed by Tukey multiple comparison test. mRNA levels are normalized against*rna18s* and *ef1a*. † Only one immature fish was available on June 7^th^. It is presented for graphical purposes only and is not included in the statistical analysis.

In under-yearling immature fish stable *fshb* mRNA levels were observed in July/August, then decreased from August 22^nd^ to October 23^rd^. Maturing fish, which were first identified on September 27^th^, had stable *fshb* transcript levels until last day of sampling on October 23^rd^. Maturing fish showed higher *fshb* mRNA levels than immature fish, from the date they were first identified to the end of the sampling period (September 27^th^ to October 23^rd^; Fig 6B). Both maturing and immature fish show stable *lhb* mRNA levels over time between July 4^th^ and October 23^rd^. No differences in *lhb* mRNA abundance are detected between groups at any time point (Fig 6D).

### 3.4 Androgens plasma levels

Plasma levels of T and 11-KT were measured only in one year old fish. T levels of one-year old maturing parr increased over time from 0.74 ± 0.07 ng/mL on April 25^th^, to 1.72 ± 0.16 ng/mL on July 4^th^, while remaining stable between 0.2 to 0.3 ng/mL in immature fish during the same period. Maturing fish showed statistically significant higher T plasma level than immature fish on May 8^th^ and 23^rd^ (Fig 7A). In contrast, 11-KT plasma levels remained below the detection limit (0.1 ng/mL) throughout the season (Fig 7B) in immature fish and increased from non-detectable (below 0.1 ng/mL) in almost all maturing males on April 25^th^ to 2.07 ± 0,57 ng/mL on July 4^th^.

**Figure 7.**
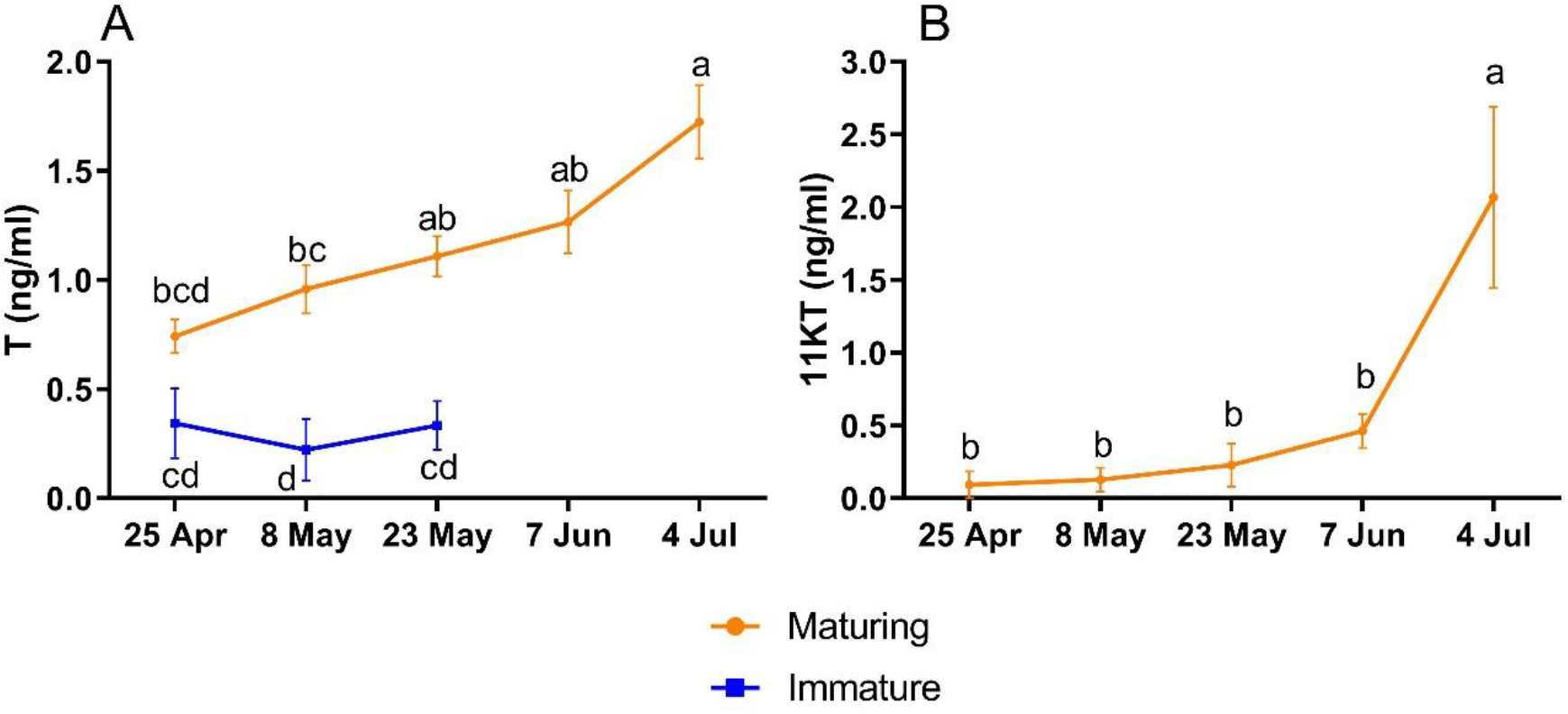
**A)** Testosterone (T) and **B)** 11-ketotestosterone (11-KT) plasma levels in one-year old male parr. Data are presented as mean ± SEM. Distinct letters denote statistically significant differences among groups (p<0.05), analysed via two-way ANOVA followed by Tukey multiple comparison test. Plasma 11-KT was not detected in immature fish. The number of biological replicates per point is listed in table 1.

### 3.5 Characterization of testis developmental stages

For the subsequent analysis of changes in gonadotropins mRNA levels, GSI and steroid plasma levels, the fish were grouped according to their testis developmental stage.

A significant increase in *fshb* transcripts was detected between stage I and stage II-to-IV (2.1- to 4.8-fold increase, Fig 8A). Fish in stages II, III and VI showed significantly higher *lhb* mRNA levels compared to stage I individuals (3.7- to 7.2-fold increase Fig. 8B). Fish at stage I had average GSI < 0.05 (0.032 ± 0.004). This value increased between 0.2 to 0.4 at stages II to V and rose to 1.95 at stage VI (Fig. 8C). 11-KT plasma content remained below detection limit (0.1 ng/mL) in all (except one) stage I, II and III fish. Values ranging from 0.30 ± 0.13 ng/mL (stage IV) to 1.34 ± 0.43 ng/mL (stage VI) were detected in more advanced stages, but with no statistically significant differences between groups, likely due to the high variation between individuals (Fig. 8D). Plasma T levels, on the other hand, were detectable in some fish from stage I (0.27 ± 0.12 ng/mL), steadily increasing up to 1.43 ± 0.12 ng/mL at stage VI (Fig. 8E).

**Figure 8.**
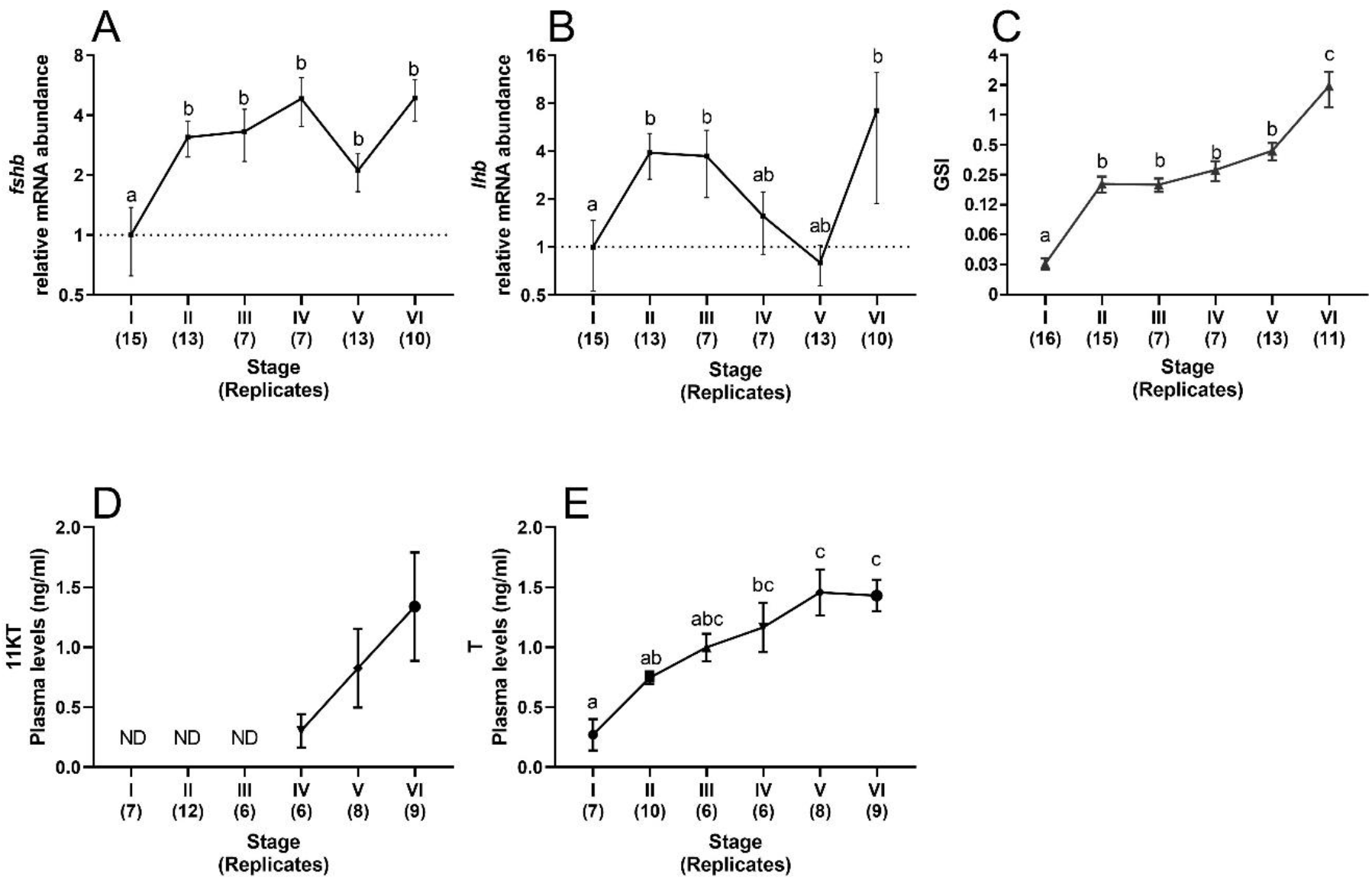
Changes in **A)** *fshb*, **B)** *lhb*, **C)** GSI, **D)** 11-ketotestosterone (11KT), **E)** testosterone (T), according to testis developmental stages I-VI. The transcript abundance (*fshb*, *lhb*) of stage I fish was set as value 1 for the representation and the following stages are represented as fold-induction. mRNA levels were normalized against *rna18s* and *ef1a*. Data are presented as mean ± SEM from pooled under-yearling and one-year old fish. Distinct letters denote statistically significant differences among groups (p<0.05), analysed via one-way ANOVA, followed by Tukey multiple comparison test. The number of available biological replicates available for each variable is shown below each stage. ND; not detected.

## 4 Discussion

This study investigates timing and age of sexual maturation in Atlantic salmon male parr in a farmed second generation from wild caught from the river Figgjo (South-west Norway), reared under natural photoperiod and water temperature.

Atlantic salmon spawns in autumn/winter with gonadal maturation in male parr occurring months earlier, generally in spring (Aas et al., 2011). In the present study, one year old male parr showed first sign of sexual maturation in spring, starting before the end of April. This is earlier than what has been previously reported. For instance, (Maugars and Schmitz, 2008) reported the first signs of maturation in male parr in early June, and Mayer et al. (1990) in July. In these studies, the fish, belonging to different strains, were reared at a higher latitude compared to our study (63° N vs 58° N), thus experiencing different environmental conditions. The variability in the maturation period between different strains, breeding in different rivers, is a known characteristic of Atlantic salmon, aimed at synchronising reproductive behaviour with the surrounding environment (Aas et al., 2011). However, despite the numerous studies investigating the link between environment, genetic background and sexual maturation, a detailed understanding of the molecular mechanism involved in the regulation of puberty in Atlantic salmon is still missing.

The levels of gonadotropin mRNA found in the one-year old males were also peculiar, as no statistically significant differences were detected between maturing and immature fish (except on May 23^rd^), despite a tendency for higher abundance in maturing males. In salmonids, both *fshb* mRNA and Fsh protein plasma levels increase during the early stages of maturation, at the onset of spermatogenesis, while *lhb* mRNA and Lh plasma protein levels are very low or undetectable at this early stages, increasing significantly toward the spawning season (Campbell et al., 2003; Gomez et al., 1999; Planas and Swanson, 1995; Swanson et al., 1991). It is possible that biologically relevant differences might have been too small to be statistically significant, or that a peak in gonadotropin mRNA levels had occurred outside the sampling period. On the other hand, plasma T and 11-KT concentrations increased gradually in maturing males while remaining at stable lower levels (T) or undetectable (11-KT) in immature males, in concert with their role in sexual maturation (Fontaine et al., 2020b; Schulz et al., 2010). While androgen plasma concentrations are comparable with previous studies (Maugars and Schmitz, 2008; Mayer et al., 1990a; Stuart‐Kregor et al., 1981), the androgen increase observed in maturing males occurred 1-2 months earlier in the year than what has been reported previously, but consistent with the observed increase in GSI. Histological analyses confirmed the advancement in testis development in fish with GSI > 0.05 (ranging from stage II to VI) and the presence of only spermatogonia type A (SPA; stage I) in the testis of fish with GSI ≤ 0.05. In the present study, testis exhibiting either, or both, undifferentiated (SPA_und_) and differentiated (SPA_diff_) SPA cysts were included in stage I. However, a number of recent studies (Crespo et al., 2019; Kjærner-Semb et al., 2018; Melo et al., 2014; Schulz et al., 2019; Skaftnesmo et al., 2017) have shown that detecting and quantifying proliferating SPA allows identification of males who have initiated sexual maturation even at this early stage. It is therefore possible that some of our stage I samples include fish that indeed have entered the maturation process. Such analysis was not performed here as residual spermatozoa were detected in a subset of the one-year old maturing males, thus suggesting that for some fish sexual maturation had been initiated for the first time at a younger age, and that these fish were maturing for the second time at this stage. To determine the age of first maturation for this subgroup, we sampled under-yearling fish next.

Under-yearling males showed the first signs of sexual maturation in autumn, as early as six months after hatching. As expected, maturing under-yearlings, initially grouped by a GSI > 0.05, displayed a 25- to 115-fold induction in pituitary *fshb* mRNA levels, but not *lhb* mRNA transcripts, compared to immature fish. As previously mentioned, in salmonids, *fshb* mRNA transcripts peak during early stages of maturation, while *lhb* mRNA are very low or undetectable, rising instead toward the spawning season and sperm maturation (Campbell et al., 2003; Gomez et al., 1999; Planas and Swanson, 1995; Swanson et al., 1991). Concomitant with the increase in *fshb* mRNA level, histological analyses confirmed the advancement in testis development, ranging from stage II to V in September. Maturing fish reached stage VI, characterised by the predominant presence of spermatozoa in the testes, in October. Such advanced testis development suggests that it is unlikely that these fish were undergoing a “dummy run”, which is characterized by an incomplete activation of the BPG-axis (Okuzawa, 2002). Interestingly, maturing males reached the final stages of testis development in autumn, during the natural spawning season in Scandinavian rivers (Aas et al., 2011). In the wild, sexually maturing under-yearling male parr have been identified (by milt production) in Atlantic salmon in a number of rivers in the Armorican massif, in northwest France, most likely due to the higher temperatures found in this system (Bagliniere and Maisse, 1985). While under-yearling maturation could be induced by photoperiod manipulation (LD 12:12 followed by LD 24:0) in a population of farmed Atlantic salmon in western Norway (Nordgarden et al., 2007), the present study now demonstrates that under-yearling sexual maturation can also occur under standard farming condition, using natural photoperiod and water temperature.

For further characterisation of the physiological conditions of male parr during development, gonadotropin mRNA levels, GSI and steroid plasma levels were measured between the different testis developmental stages, pooling both age groups together. A statistically significant 2- to 6-fold increase in pituitary *fshb* transcripts was detected between stage I and the more advanced stages (II-VI). This is a lower induction compared to other studies conducted on maturing parr (Maugars and Schmitz, 2008) and smolt (Melo et al., 2014), showing a 20- to 60- and 100- to 200-fold increase respectively. Similarly, the relative abundance of *lhb* transcripts detected in the present study showed an 8-fold induction between stage I and VI while increases of 100- or 600-fold was reported in the aforementioned studies. While circulating steroids levels also increased from early to late developmental stages, the measured concentrations were generally lower compared to previous studies. In particular 11-KT levels remained below detection limit in most fish up to developmental stage III, when spermatogonia (type A and B) and spermatocytes are already present in the testes. This was unexpected, as 11-KT is involved in germ cell maturation and proliferation (Schulz et al., 2010). However, the measured plasma 11-KT acting as endocrine messenger to target organs, may differ from *in situ* levels in the testes acting as paracrine signal to promote germ cell maturation. Despite the values of the investigated parameters being lower than expected, testis development proceeded up to later stages of maturation. The comparison of our results with the literature indicates a high level of variability between studies. Our results also suggest that small variations in gonadotropin transcripts and androgen plasma levels might suffice to promote sexual maturation.

Histological analyses revealed the presence of early previtellogenic oocytes in 8.7 % of testes analysed, including both under-yearling and one-year old fish. The intersex condition has been documented in different gonochoristic (fixed-sex) teleost species in both wild and laboratory animals (Bahamonde et al., 2013) and is considered as a characteristic effect of exposure to endocrine disrupting compounds, most commonly estrogenic chemicals (Metcalfe et al., 2010). In male rainbow trout (*Oncorhynchus mykiss*), exposure of estrogenic compounds elicit a dose-dependent synthesis of vitellogenin and a concomitant inhibition of testicular growth (Jobling et al., 1996) and gonadal intersex (Depiereux et al., 2014). Intersex as a natural phenomenon in the wild is very challenging to evaluate given the difficulty to find pristine, uncontaminated reference sites but also due to the influence of the season of sampling age and maturity state of the animal (Bahamonde et al., 2013). The available data suggest that in salmonids, the disorder has a very low occurrence as spontaneous intersex has been reported in only 1 individual over 2660 chinook salmon (*Oncorhyncus tshawytscha*) sampled over 3 years in New Zealand and 3000 coho salmon (*Oncorhyncus. kisutch*) in Chile (Kinnison et al., 2000). The low natural occurrence of intersex in salmonids suggests that the feminization reported in the present study could be indicative of contamination from endocrine-disruptive compounds in the water. However, in those testes exhibiting ovotestis characteristics, there was only a few scattered oocytes present in the tissue suggesting the occurrence of only a mild estrogenic effect.

## 5 Conclusion

The present work represents an investigation on early sexual maturation in Atlantic salmon male parr. The timing and characterization of testes development were determined by measuring key morphometric and physiological traits in both maturing and non-maturing male parr over two consecutive seasons. These traits, established indicators of sexual maturation, included morphometric parameters [body weight, fork length, gonad weight, gonadosomatic index (GSI) and condition factor (K)], plasma steroids levels (T, 11-KT), gonadotropin gene expression levels (*fshb*, *lhb*) and testes developmental stages. The analyses revealed that male parr can mature in Autumn, as early as six months after hatching. In spring, at one year of age, gonadal maturation is activated, including both fish that matured as under-yearlings and first-time maturing parr. This study highlights the importance of histological analyses when investigating pubertal activation in salmonids. This is the first study comparing the physiology of under-yearling and one-year old maturing male parr, and further demonstrates the remarkable male reproductive plasticity displayed by Atlantic salmon.

## Supporting information

Photoperiod

Water temperature

Testes histology. Incomplete maturation

## Acknowledgements

The authors wish to thank the staff at the NINA Aquatic Research Station at Ims, in particular Mr. Knut Aanestad Bergersen, for providing and maintaining the fish. We are grateful to Dr. Gersende Maugars for her revision of the manuscript.

## Fundings

This project has received funding from the European Union’s Horizon 2020 research and innovation programme under the Marie Skłodowska-Curie grant agreement No 642893 (IMPRESS) and from the Norwegian University of Life Sciences (NMBU).

## Declaration of interest

The authors declare that they have no known competing financial interests or personal relationships that could have appeared to influence the work reported in this paper.

## Authors contribution

**EC**: Conceptualization, Methodology, formal analysis, Investigation, Writing – Original Draft, Visualization. **KvK**: Investigation, Writing – Review & Editing, Visualization. **RNL**: Investigation, Methodology. **IM**: Investigation, Writing – Review & Editing. **RF**: Conceptualization, Methodology, Writing – Review & Editing, Supervision. **FAW**: Conceptualization, Methodology, Resources, Writing – Review & Editing, Supervision, Project administration, Funding acquisition

## Supplementary files

**Figure S1.** Natural photoperiod at Ims, Norway (black line; 58°54’N, 5°57’E) and at the Norrfors hatchery, Sweden (red dotted line; 63°N, 20°E) where the studies from (Maugars and Schmitz, 2008; Mayer et al., 1990b) were performed. Y axes indicates time of the day. Sunrise and sunset time represented from lower and upper line respectively. Data were retrieved from https://www.timeanddate.com

**Figure S2.** Seasonal changes in water temperature at Ims research station (black line; 58°54’N, 5°57’E) from 2015 to 2017. Water temperature at Norrfors hatchery (red dotted line; 63°N,20°E) in 2003 where the studies from (Maugars and Schmitz, 2008; Mayer et al., 1990b) was performed. Temperature data from Norrfors hatchery were kindly provided from Dr. Gersende Maugars.

**Figure S3.** Sagittal section showing incomplete maturation of salmon testis, where only parts of the tissue is maturing. Sections (3 μm) were prepared in plastic resin and stained with Toluidine Blue O. Scale bar 100 μm.

## Notes

### Competing Interest Statement

The authors have declared no competing interest.

