## Supplementary figures and images for "Sexual maturation in Atlantic salmon male parr may be triggered both in early spring and late summer under standard farming conditions"

### Fig S1 .tif

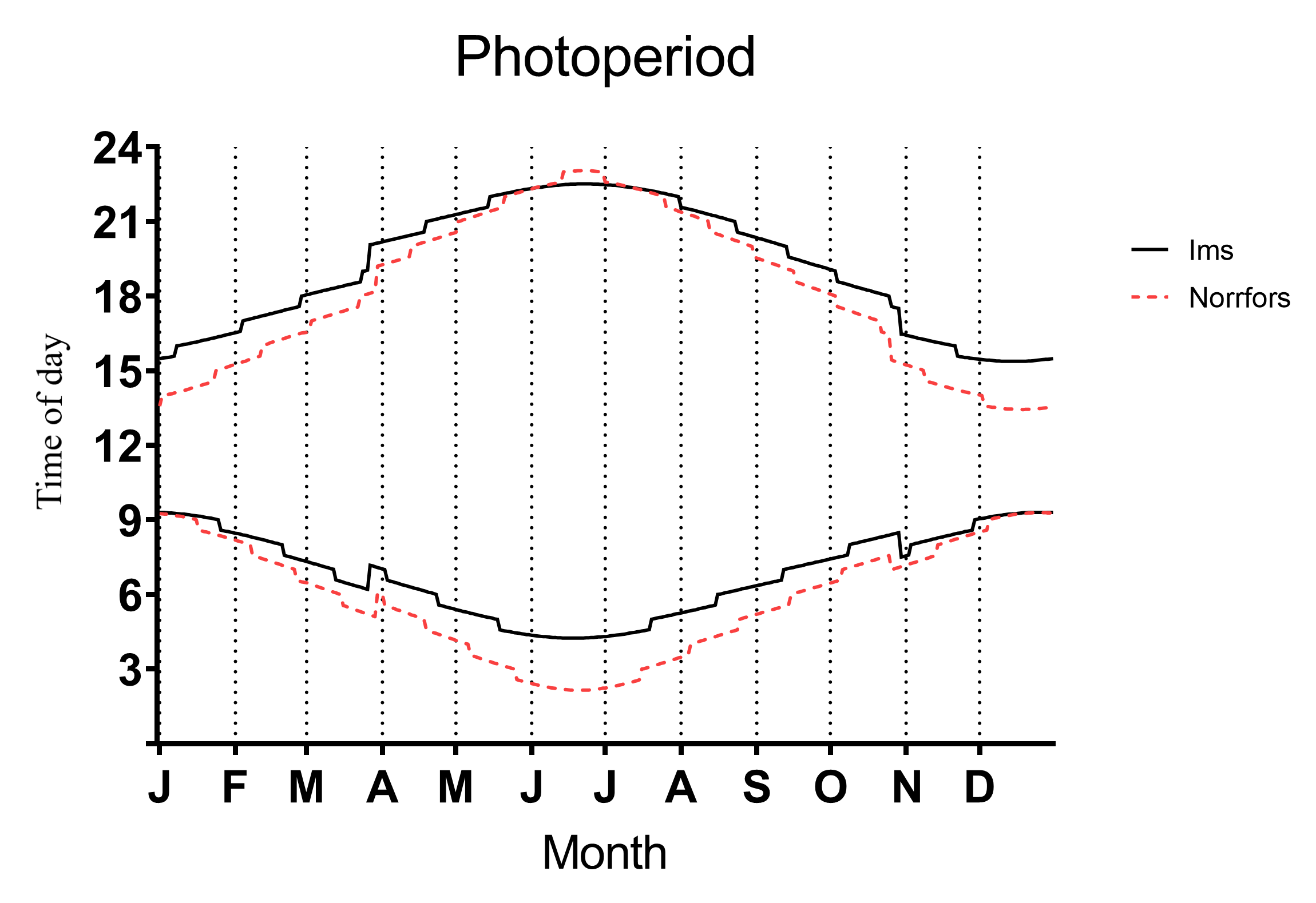

### Fig S2 .tif

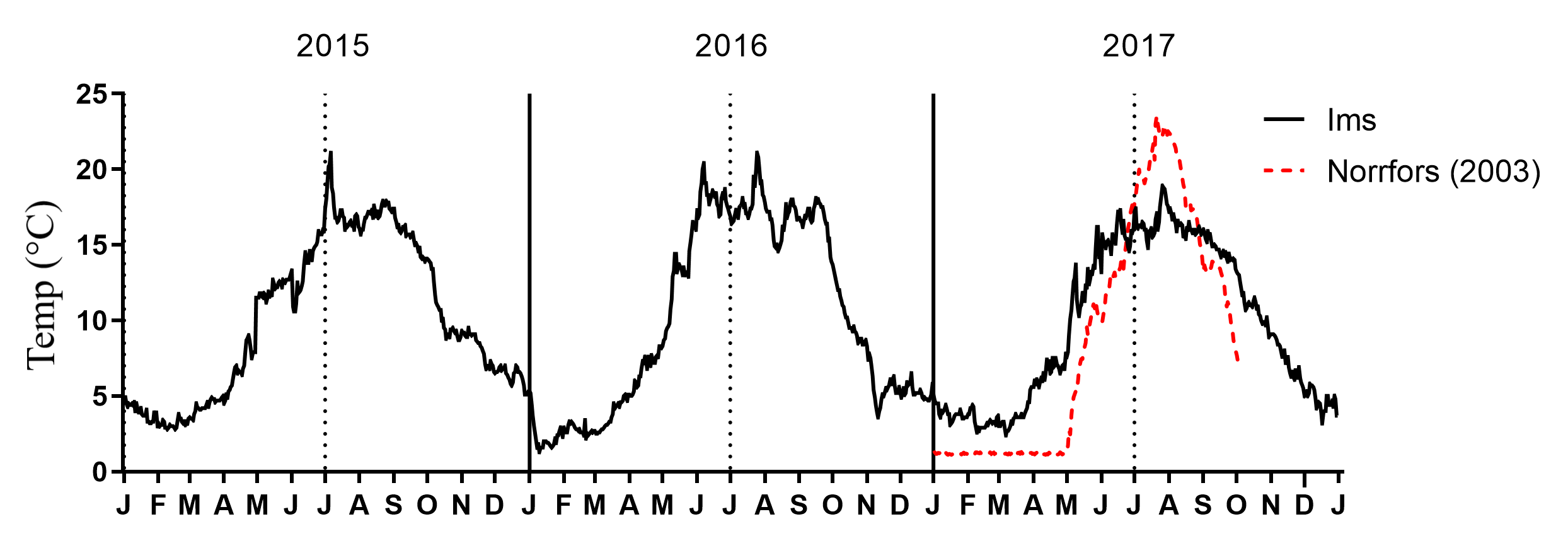

### Fig S3.tif

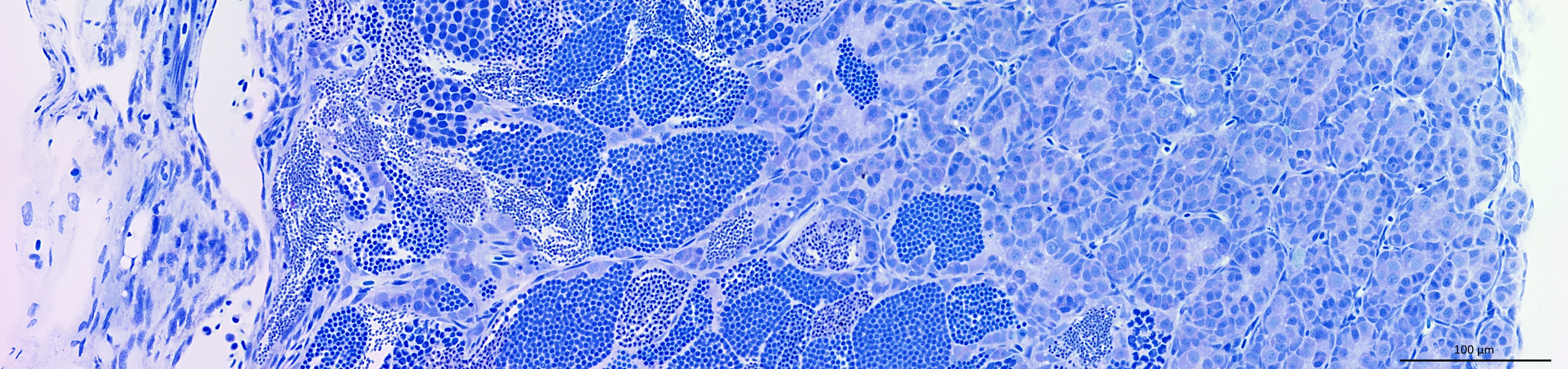
